# Benefits of living closer to kin vary by genealogical relationship in a territorial mammal

**DOI:** 10.1101/2021.11.30.470584

**Authors:** Sam F. Walmsley, Stan Boutin, Ben Dantzer, Jeffrey E. Lane, David W. Coltman, Andrew G. McAdam

## Abstract

While cooperative interactions among kin are a key building block in the societies of group-living species, their importance for species with more variable social environments is unclear. North American red squirrels (*Tamiasciurus hudsonicus*) defend individual territories in dynamic neighbourhoods and are known to benefit from living among familiar conspecifics, but not relatives. However, kin-directed behaviours may be restricted to specific genealogical relationships or strongly mediated by geographic distance, masking their influence at broader scales. Using distance between territories as a proxy for the ability of individuals to interact, we estimated the influence of primary kin (parents, offspring, siblings) on the annual survival and reproductive success of red squirrels. This approach revealed associations between fitness and access to kin, but only for certain genealogical relationships and fitness components. For example, females had enhanced annual survival when living closer to their daughters, though the reverse was not true. Most surprising was the finding that males had higher annual reproductive success when living closer to their father, suggesting possible recognition and cooperation among fathers and sons. Together, these findings point to unexpected nuance in the fitness consequences of kinship dynamics for a species that is territorial and largely solitary.

## Introduction

How individuals disperse, settle, and die has important consequences for the evolution of cooperation, constraining the mechanisms by which affiliative relationships can form [1,2]. For example, in most group-living mammals, female-biased philopatry results in social units composed of matrilineal relatives [3]. Here, kin selection provides the opportunity for the development of stable and cooperative groups, bolstered by the additional benefits of inclusive fitness [4,3]. While the challenges of navigating group dynamics can be significant, members often rely on predictable dominance and kinship-based hierarchies to structure their interactions. For example, spotted hyenas (*Crocuta crocuta*) are born into matrilineal clans and inherit the social rank and network of their mother, a key determinant of lifelong fitness [5–7]. Less understood is whether kinship shapes cooperative relationships in species where access to kin is more variable [8–10], especially those lacking obvious social units, such as solitary species [11].

Non-traditional cooperative relationships among kin may be particularly difficult to study if they rely on more cryptic forms of interaction. For example, kin-directed behaviours may simply involve increased spatial tolerance (e.g., Galápagos sea lions, *Zalophus wollebaeki* [12]; Atlantic salmon, *Salmo salar* [13]), or subtle reductions in antagonism (e.g., gray squirrels, *Sciurus carolinensis* [14]). An important follow-up question is whether these subtler processes should translate into meaningful effects on individual variation in survival and reproductive success. Quantifying the fitness consequences of these less-conspicuous interactions is a crucial step towards understanding whether the benefits of living with kin are mirrored in species with highly variable social environments.

Evidence from group-living species suggests that variable access to kin can result in a compensatory investment in mutualistic relationships with non-relatives. Vampire bats (*Desmodus rotundus*) rely on regular food sharing, which typically occurs among close kin [15]. However, to mitigate the risk of resting at a different roost from these preferred partners, individuals “hedge their bets” by simultaneously developing affiliative relationships with unrelated conspecifics. Similar strategies are found in baboons: female yellow baboons (*Papio cynocephalus*) preferentially associate with their mothers or daughters but rely on bonds with non-kin when necessary [16], and female Chacma baboons (*Papio ursinus*) increase investment in new grooming relationships when a preferred kin partner dies [17]. These findings demonstrate that cooperation with unrelated individuals can buffer individuals from the risks of temporarily or permanently losing access to a close relative. In other words, despite lacking the advantage of inclusive fitness benefits, group-living animals may choose to invest in mutualistic bonds with non-kin if it leads to a more predictable social environment [18,19]. So, while kin selection is typically considered the lower-threshold pathway to prosocial bonds [20], mammals living in less-structured social environments may instead prioritise mutualistic bonds with potentially unrelated individuals.

North American red squirrels (*Tamiasciurus hudsonicus*; hereafter red squirrels) defend exclusive territories containing a central hoard of stored food resources (midden) from nearby conspecifics. Populations are characterized by fluctuating membership, driven by an influx of juveniles in the summertime [21], particularly during episodic resource pulses. Red squirrels sometimes establish territories near their parents [21,22], creating some spatial scaffolding for kin cooperation. However, mean relatedness among neighbours remains low [23], and future access to kin depends almost entirely on where an individual can establish a territory. Compared to many group-living mammals, red squirrels cannot count on living near relatives as adults. So, are red squirrels exploiting this variation in access to kin, or does the unpredictability of the social environment lead to a prioritization of mutualistic bonds with available neighbours, regardless of relatedness?

Siracusa et al. (2021) found that familiarity, defined as the amount of time two individuals lived within 130 m of one another, enhanced the survival and reproductive success of red squirrels whereas the mean relatedness of neighbouring conspecifics had no effect [23]. Though consistent with the hypothesis that animals benefit by social bet-hedging when access to kin is unpredictable [15], this finding is surprising given evidence of rare but seemingly cooperative kin-biased behaviour. Related red squirrels sometimes nest communally for warmth [24], adopt orphaned juveniles [25], and even bequeath entire territories to their offspring [21]. Possible explanations for this discord could be that benefits of kin-directed behaviour only occur at finer spatial scales [23], or perhaps, that kin benefits are limited to specific genealogical relationships (e.g., mother-daughter) [26,27], especially if kin recognition and/or affiliation are contingent on early associations in the natal nest [28], which may be the case for red squirrels [29].

Leveraging a long-term study of wild red squirrels in Yukon, Canada, we ask whether kinship plays a role in structuring fitness-enhancing interactions in a social environment where access to kin is highly variable across individuals and over time. Red squirrels risk intrusions from neighbours when away from their territory [30], likely making it costly to interact with individuals that are far away. Following studies on kin benefits in humans [31], we aimed to exploit the spatial limitations of any affiliative behaviours, hypothesizing that the influence of kin on annualized survival and reproductive success would be mediated by geographic distance. Given the possibility that beneficial kin-directed behaviours vary by genealogical relationship [26,32], we focused our analysis on primary kin (mothers, fathers, sons, daughters, full siblings), estimating separate influences for each kinship class. We predicted that any influences of kin on annual survival or annual reproductive success would dissipate as a function of distance. And while we were interested in possible interactions among fathers and offspring, we expected that benefits were more likely to occur among kin that shared a natal nest (e.g., mother-offspring dyads).

## Materials and methods

### Study site and long-term data collection

All data used in this study were collected as part of the Kluane Red Squirrel Project (KRSP), a long-term research project investigating populations of wild red squirrels living in southwestern Yukon, Canada (61°N, 138°W). Since 1987, KRSP have tagged, enumerated, and monitored individual red squirrels throughout their lives using alphanumeric ear tags (Monel #1; National Band and Tag, Newport, KY, USA). Red squirrel territories are typically centred on caches (middens) containing hoarded white spruce cones (*Picea glauca*). White spruce trees undergo masting events every 4-7 years, resulting in resource pulses for red squirrels. Our analysis focused on two study areas (“Kloo” and “Sulphur”). Both study areas were staked at 30 m intervals, with locations identified to tenths of an interval, meaning that any spatial position could be specified within approximately 3 m. Both male and female red squirrels are exclusively territorial, and typically defend the same territory year-round for life, allowing us to reliably locate individuals and map out the distances between them. Territory sizes vary considerably in this population [33], but are estimated to be 0.34 ha on average [34], corresponding to a radius of approximately 33 m for a hypothetical circular territory. Territory ownership was determined during a census each May through repeated live-trapping at each midden or nest on the study area (Tomahawk Live Trap Co., Tomahawk, WI, USA), and by behavioural observations (principally, evidence that an individual consistently used its territorial “rattle” vocalization at a given location [30]). Pregnant females are regularly monitored in the spring so that juveniles can be tagged and linked to their maternal pedigree. Previous work has confirmed that these linkages are unlikely to be biased by rare adoption events [25]. Starting in 2003, all censused individuals have also been genotyped at 16 microsatellite loci. These data have been used to supplement maternal linkages with paternities (99% confidence, CERVUS 3.0 [35]), resulting in a comprehensive, long-term pedigree including nearly every individual living on each study area [36]. For additional information on the study site and long-term data collection protocols, see [21,37,38].

### Study subjects

For all known red squirrels living on the Kloo or Sulphur study areas between 2003-2014, we identified pairwise relationships among primary kin (mother, father, son, daughter, full sibling) using the multigenerational pedigree and the *pedantics* package [39]. Full sibling relationships were not partitioned by sex due to a lack of data: high rates of both multiple paternity [40], and juvenile mortality meant that it was relatively rare for full siblings to persist in the population (median 2 sibling dyads per study area per year, range 0-7). Given our interest in estimating annual survival, we only included individuals with known birth years, and excluded any squirrels that were known to have died of un-natural causes, as these deaths were expected to be unrelated to influences of the social environment. Geographic distance between kin territories was measured as the linear distance between the centre of the primary midden (cone cache) of each squirrel, which is typically situated at the centre of the territory. Red squirrel middens are easily identifiable as piles of spruce cone scales and spines, often containing tunnels where cones are stored. We included territory locations of several individuals who lacked full, established middens but were known to defend a nest in the study area (n = 4). For individuals without any primary kin in the study area in a given year, we calculated their distance to the territory of a randomly selected individual, coded as “Non-kin”. Note that this category could include more distantly related kin, but no full siblings, parents, or offspring. These “Non-kin” observations served as a reference category, anchoring estimated influences of primary kin in subsequent models to a meaningful baseline. Following Siracusa et al. [23], we also calculated familiarity for each observation as the number of days that the dyad was known to live within 130 m of one another. This measure was mean-centered and scaled to have a standard deviation of 1 prior to further analysis. The distance threshold of 130 m corresponds to the estimated range at which red squirrel territorial vocalizations are heard, i.e., the acoustic social neighbourhood [41], and also approximates the spatial scale at which red squirrels respond to variation in density [42].

Prior to exploring fitness effects, we used a generalized linear mixed model (GLMM) with Gamma family (log link) to test whether males and females differed in distance to their nearest primary kin. To avoid repeated inclusions of the same observation, we randomly sampled a single distance measure per dyad for this model. We also included a random effect to account for repeated measurements of an individual across multiple dyads.

### Estimating influences of kin on survival

We measured annual survival as the detection of an individual on or after 1 January of the following calendar year (note however that trapping typically began each year at approximately 1 March). Because red squirrels rarely change territory locations [43], individuals were assumed to have died if they disappeared from the study area and were never seen or trapped again. Previous work on this population has estimated the probability of re-detecting living adults to be 100% [44].

Prior to analysis, we subset the data to only include observations of the *nearest* parent, offspring, or full sibling for each focal-year combination, resulting in a dataset with distances to different types of kin. This allowed us to directly test whether influences on fitness differed by kin relationship, while also focusing on the smallest distance values, expecting that any behaviourally mediated influences should be most apparent when squirrels were very close to one another.

First, we used generalized additive models (GAMs) to investigate possible non-linearities in the relationships between fitness and distance to specific types of primary kin. This was necessary given that influences of other squirrels could be negligible at very large distances, in which case a linear fit would not be justifiable for the full spatial extent of the data. We fitted GAMs with a binomial family and log link to relate annualized survival as the response variable. We used a basis dimension of 3 for each GAM, based on examination of the estimated degrees of freedom after fitting [45]. Crucially, we fitted an interaction between relationship type and linear distance. This allowed us to test whether the fitness-distance relationships were significantly different between different types of primary kin, compared to the reference category (‘Non-kin’). We also included mast year (yes/no) as a predictor, expecting that resource pulses could be a common cause influencing survival, and potentially, access to primary kin (i.e., environmental co-variance). Lastly, we included quadratic effects of age given the expectation that the probability of annual survival should decrease for both juveniles and senescent adults [23].

These preliminary GAMs indicated potentially non-linear relationships between fitness and distance to specific types of primary kin. However, relationships within the previously defined “social neighbourhood” in this population [23,29], i.e., up to a maximum distance between territories of 130 m, tended to be linear, with apparent changes in the relationship typically occurring at or beyond this point. Accordingly, we fitted generalized linear mixed models (GLMMs) with linear effects of distance to the restricted, “social neighbourhood” datasets, only including distance records less than or equal to 130 m. These models were fitted with binomial family and a logit link function and used the same response and predictor structure as above, explaining survival as a function of mast year, age (linear and quadratic terms), and an interaction between genealogical relationship and geographic distance. Here, we also included dyadic measures of familiarity (following the method for calculating familiarity from Siracusa et al. 2021 [23]) so that benefits of kin were estimated over and above any influences of familiarity-driven relationships. Though these within-neighbourhood (i.e., < 130 m) influences of kin were most relevant for our research question, we also provide additional GLMMs in the supplement that include distances spanning the full range of each study area as well as a quadratic effect of distance to account for apparent non-linear associations at this scale (See electronic supplementary material).

All GLMMs were fitted with crossed random effects to account for the inclusion of multiple measures of the focal squirrel (focal ID) across years and of possibly bi-directional records of the same dyad of squirrels (dyad ID). We also included a random effect for year of the study period to account for any annual variation in fitness measures not captured by the mast-year fixed effect.

### Estimating influences of kin on reproductive success

Following previous work on this population [23], we calculated separate metrics of annual reproductive success (ARS) for males and females to reflect differences in parental care. For females, we defined annual reproductive success as the number of pups that were successfully recruited (i.e., were confirmed alive for at least 200 days), excluding any non-breeding individuals. As with measures of annual survival, we first used GAMs with a Poisson family and log link to visualize any non-linear influences of each type of primary kin on annual reproductive success.

To test the sensitivity of our results to this exclusion, we also fitted identical GAMs including non-breeding females (*n* = 181), attributing them ARS values of 0. Because male red squirrels provide no obvious paternal care, annual reproductive success for males was measured as the number of pups sired per year.

As with the annual survival models, we identified possible non-linearities across the full spatial extent of each study area, but linear relationships within the social neighbourhood. Here we used GLMMs with the Poisson family and log link function to explain reproductive success as a function of distance to primary kin, fitted with an interaction between distance and genealogical relationship type. We also included dyadic familiarity, mast year, linear and quadratic effects of age, and crossed random intercepts representing variation across individuals, dyads, and years.

The model with male reproductive success as the response variable showed moderate overdispersion, so was re-fitted using the negative binomial distribution. For all GLMMs, measures of distance were log10-transformed, reducing the leverage of very large distance values. No additional mean-centering or scaling was applied, so that main effect coefficients correspond to benefits of each type of primary kin having a separate territory at very close distances (1 m). Note though that such close distances do not occur biologically: in our study, the closest distance between two squirrels with distinct territories was 9.5 m.

Prior to analysing any data, we developed and publicly uploaded an analysis plan to avoid inadvertently hypothesizing after results were known (“HARKing”) [46]. This plan is available at https://github.com/KluaneRedSquirrelProject/Analysis-Plans. Some changes to our approach were made during analysis, such as the inclusion of dyadic familiarity and year-specific random effects. All analyses were conducted in R version 4.0.5 [47]. GAMs were fit using the *mgcv* package while GLMMs were fit using *glmmTMB* [48,49]. For each GLMM, we used *DHARMa* to assess model diagnostics [50], and *MuMIn* to calculate conditional and marginal R^2^ [51,52]. Detailed sample sizes including the number of observations, focal individuals, and unique dyads are available in the electronic supplement (tables S1 through S9). Coding scripts are available at github.com/KluaneRedSquirrelProject/Walmsley-Projects.

## Results

In total, our main analysis included observations of annual reproductive success and overwinter survival from 679 individuals between 2003 and 2014. Compared to females, male squirrels generally lived farther from their nearest primary kin (95% CI for males: 124 – 160 m, for females: 89 – 111 m; β = 0.20 ± 0.04 s.e., p < 0.001; table S1; figure S1). As expected, the individuals in our study had highly variable access to primary kin. In any given year, it was common for both female and male red squirrels not to have any close relatives in their social neighbourhood (mean 0.56 primary kin within 130 m, range 0-6). These spatial associations were also transient, with kin dyads remaining in the same social environment for just 1.4 years on average (range 1-4; for 130 m neighbourhood). Access to kin was not much improved when considering the larger spatial scale of each study area (approx. 40 ha), with each squirrel having a mean of 0.83 primary kin in the study area (range 0-7) in a given year, and with overlap among kin lasting 1.5 years on average (range 1-4). The number of kin relationships appeared to vary in tandem with changes in population density attributable to resource pulses (see figure 1; electronic supplementary material, figure S2). Specifically, both density and the number of kin relationships tended to increase immediately after mast years, gradually declining thereafter.

**Figure 1.**
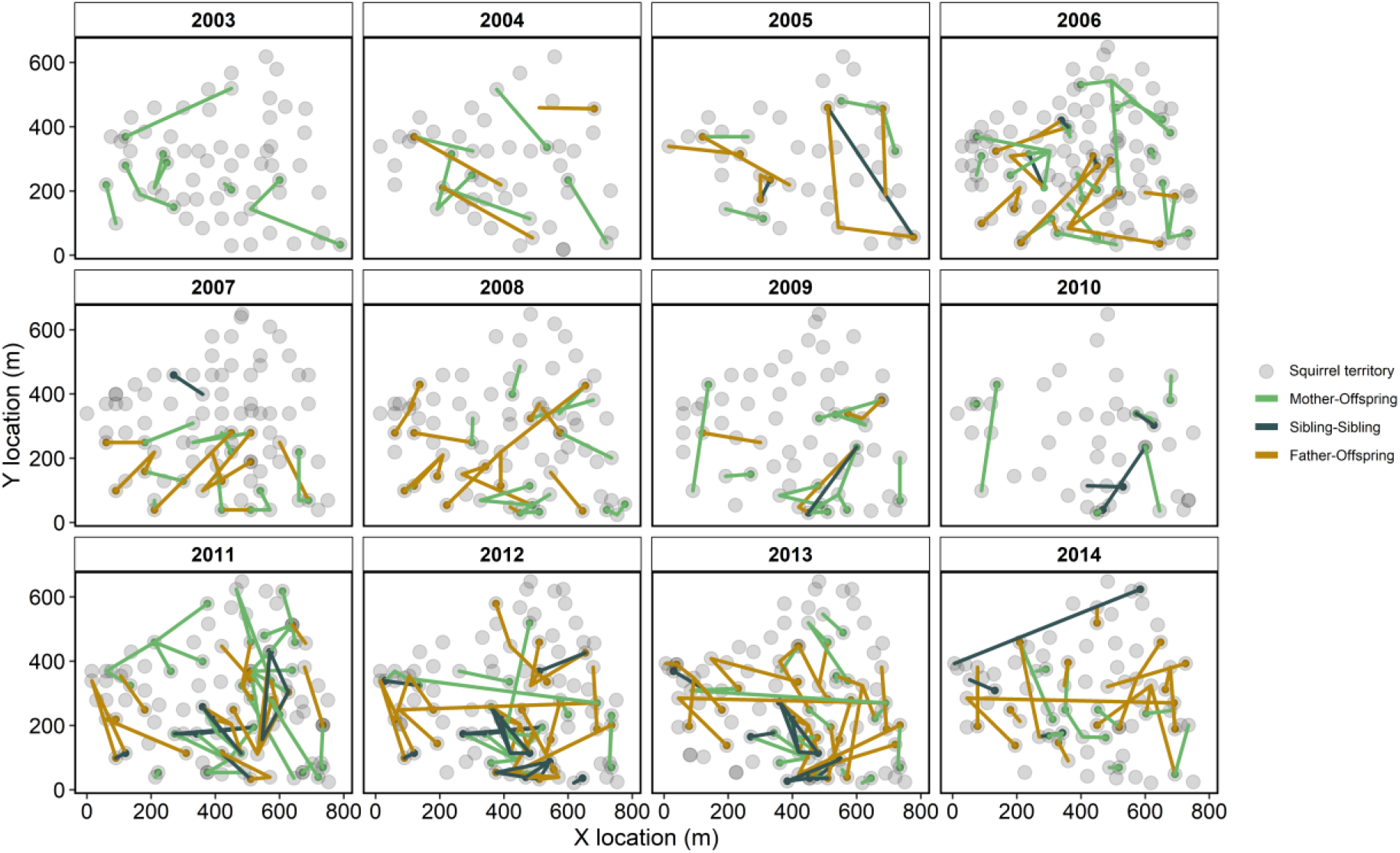
Spatial overview of kin relationships on “Kloo” study area (see figure S2 for overview of “Sulphur” area). Lines linking certain individuals represent relationships among primary kin, with colors distinguishing dyads as a sibling pair or by the sex of the parent. Grey circles show the locations of middens or territory centres for all territories occupied each year. Mast years (resource pulses) occurred in 2005, 2010, and 2014.

### Influences of kin on survival

For females, annual survival was strongly associated with access to daughters. At the scale of the social neighbourhood, females whose nearest primary kin was a daughter were more likely to survive when that daughter was closer, as evidenced by the significantly more negative distance-survival relationship for daughters compared to the reference category, “non-kin” (β = −3.78 ± 1.47 s.e., p = 0.010; table S2). Crucially, this effect waned near the threshold of the social neighbourhood (130 m), meaning that the survival benefit of having a daughter as nearest kin occurred only at short distances (e.g., less than approximately 40 m; figure 2). This effect was estimated while controlling for a quadratic effect of the age of the focal individual (i.e., in the present example, the mother), which, as expected, was also a significant predictor of survival (β^2^ = −0.10 ± 0.04 s.e., p = 0.015). The additional GLMM including distance measures greater than 130 m did not identify a significant effect of the proximity of daughters on female survival (table S3), further supporting the idea that daughter-induced benefits are only present within the finer spatial scale of the social neighbourhood.

**Figure 2.**
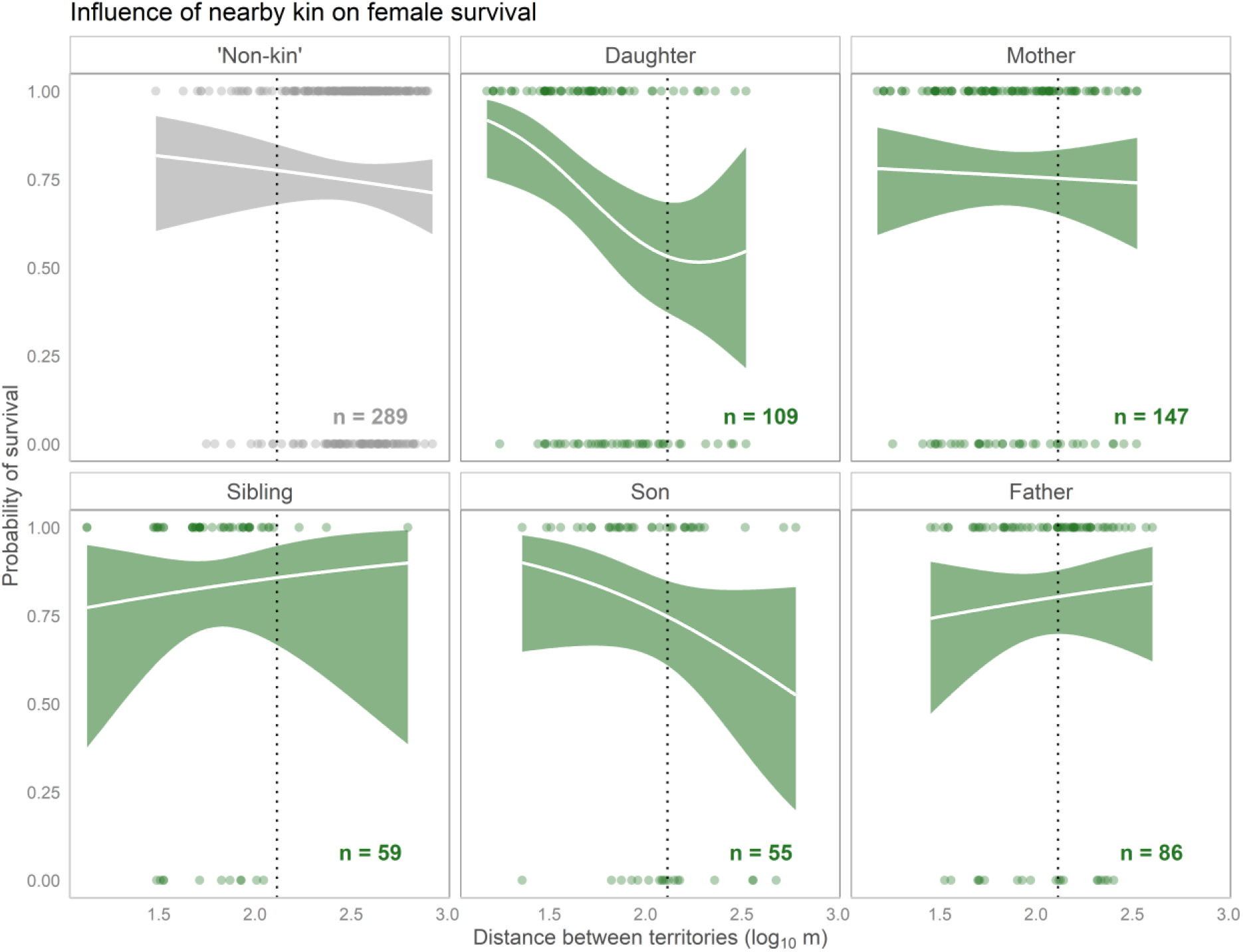
Lines represent predicted probabilities of female survival, estimated with GAMs while controlling for the influences of age (linear and quadratic) and resource pulses. Here, probabilities are shown for a focal individual of age 2 in a non-mast year. Points represent raw data, so are not an exact reflection of the partial smooth. Shaded bands show 95% confidence intervals. The dotted vertical line corresponds to 130m, a commonly used radius of red squirrels’ social neighbourhoods. ‘Non-kin’ records are made up of individuals with no primary kin in the study area, for which distances were calculated to a randomly chosen individual.

Adult males were less likely to survive during mast years compared to non-mast years (β = −0.69 ± 0.33 s.e., p = 0.036). However, there were no significant differences in the distance-survival relationship for any types of primary kin, compared to the reference effect calculated for non-kin conspecifics (table S4) indicating no effect of kin proximity on male survival. Results were similar when including the entire study area, however the quadratic component of the daughter-distance interaction was statistically significant and negative (β = −30.31 ± 11.84 s.e., p = 0.010; table S5), suggesting that male survival increases with distance to daughters up to approximately 130 m, then decreases (figure S5). For models of both male and female survival, effects of kin were identified while conditioning on familiarity between the focal and target individuals, which itself had little influence on annual survival (table S2, table S4).

### Influences of kin on reproductive success

For females, annual reproductive success was strongly influenced by masting events, with breeding females producing approximately 3 times as many recruiting offspring during mast years vs. non-mast years, on average (β = 1.16 ± 0.18 s.e., p < 0.001). However, reproductive success was not associated with distance to any type of primary kin, relative to non-kin (table S6; electronic supplementary material, figure S3). These results were similar for models including distances to kin across the entire study area (table S7). The additional analysis including non-breeding individuals (attributing them 0 pups recruited) revealed the same pattern whereby we found no evidence that distance to kin affected female reproductive success (figure S4), so we proceeded to interpret values from GLMMs using ARS of breeders alone. As with female survival, we found little influence of dyadic familiarity on fitness (β = 0.04 ± 0.06 s.e., p = 0.472).

In contrast, we found clear links between annual reproductive success and distance to kin for males. Outputs from the GAMs pointed to the possibility of sex-specific trends: males sharing their social neighbourhood with a father or son seemed to sire more pups when living closer to that individual (figure 3). Oppositely, distance to a mother or daughter’s territory seemed to be positively associated with male reproductive success (i.e., implying a relative cost of living closer to female kin; figure 3). This general pattern was supported by the parametric outputs from the GLMM, though only effects of fathers and mothers were statistically significant in the social neighbourhood analysis. For males whose nearest primary kin was their father, those living closer to their father sired more pups, reflected in the more negative distance-ARS relationship for fathers compared to non-kin (β = −5.07 ± 2.26 s.e., p = 0.025). It is worth noting that this effect was detected with a smaller sample size than some other kin-specific effects, as it was less common for males to have a father as their nearest primary kin than a mother or daughter, for example (figure 3). Oppositely, the GLMM suggested that among males whose nearest primary kin was their mother, those living *farther* from their mother were more successful (β = 2.79 ± 1.42 s.e., p = 0.049), though this effect was smaller in magnitude than the benefits of access to fathers. As before, these effects were estimated while conditioning on masting events and age (linear and quadratic terms), all of which also had statistically significant effects on male ARS (table S8). Here we also detected a positive effect of dyadic familiarity on male reproductive success (β = 0.12 ± 0.05 s.e., p = 0.022). Though distance to daughters did not significantly influence male ARS at the scale of the social neighbourhood, the effect of daughters emerged as a significant influence on the number of pups sired in the model based on the entire study area (table S9). This suggests that any influence of daughters on male reproductive success (negative but non-significant at the scale of the social neighbourhood; Table S8) likely changes after a certain distance threshold, and that nearby daughters may negatively impact their father’s reproductive success (figure 3). Finally, we detected a significant effect of distance to non-kin (table S9), suggesting that when considering a large spatial scale, males may sire more pups at higher densities overall, possibly a result of an increased availability of females. However, a similar effect was not detected when considering the social environment alone (i.e., the 130 m social neighbourhood; table S8).

**Figure 3.**
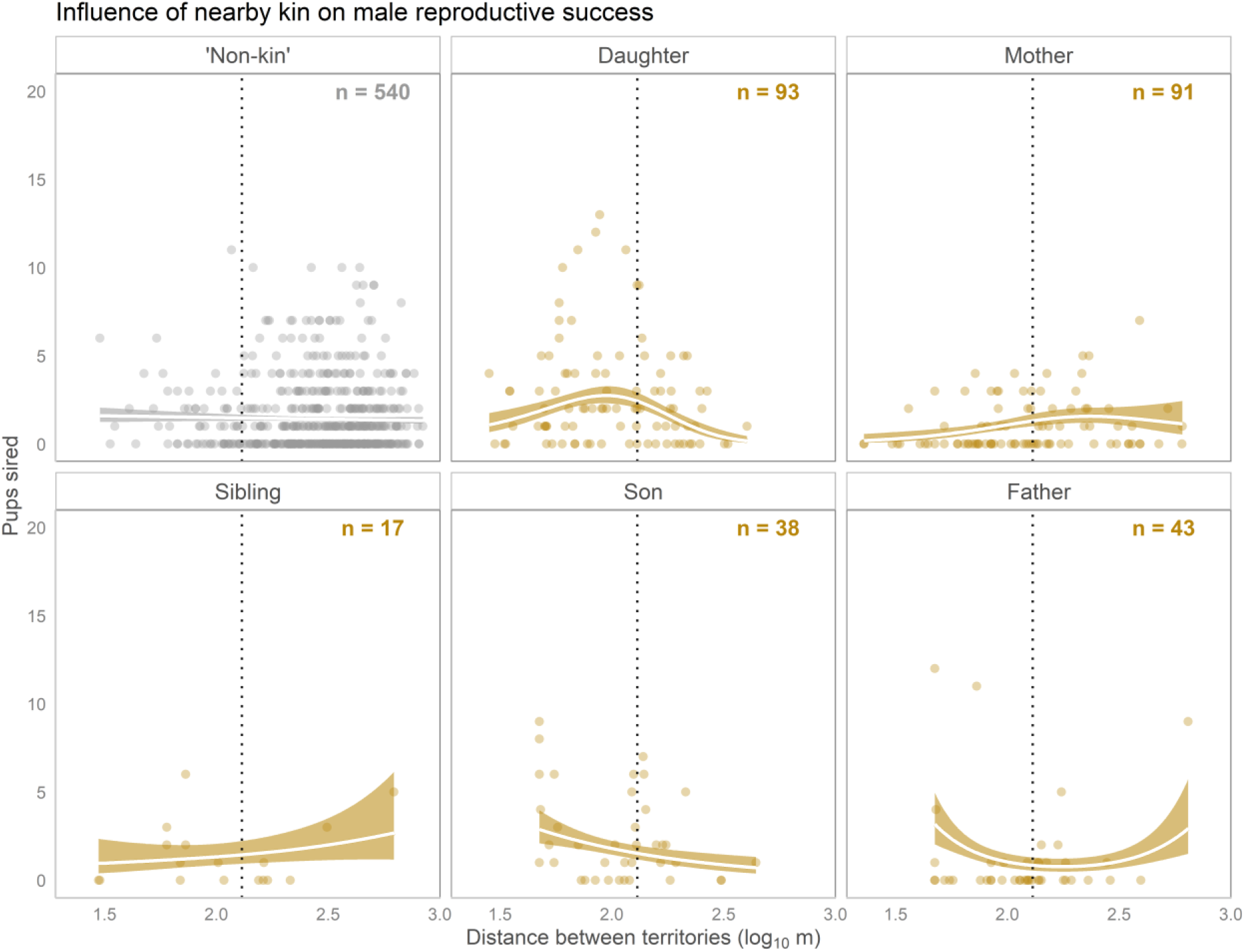
Lines represent predicted probabilities of male annual reproductive success, estimated with GAMs, controlling for the influences of age (linear and quadratic) and resource pulses, with shaded bands representing 95% confidence intervals. Here, probabilities are shown for a focal individual of age 2 in a non-mast year. Points represent raw data, so are not an exact reflection of the partial smooth. The dotted vertical line corresponds to 130 m, a commonly used radius of red squirrels’ social neighbourhoods. Non-kin records are made up of individuals with no primary kin in the study area, and distances were calculated relative to a randomly chosen individual.

## Discussion

In this study, we used the geographic distance between territories of solitary red squirrels to test whether access to kin influenced individual fitness. We found that red squirrels had improved survival and reproductive success when living near primary kin, though these effects were surprisingly nuanced, hinging on genealogical relationship, sex, and fitness measure. The dilution of fitness consequences from nearby kin across relatively short distances (0-130 m) is consistent with the hypothesis that these benefits are caused by kin-specific behavioural interactions, though we cannot rule out the possibility that other (unknown) factors also play a role. Together, our findings suggest complex kinship effects on survival and reproductive success in a solitary mammal, highlighting the value of multigenerational datasets for uncovering otherwise cryptic social dynamics.

The increased survival of mothers living closer to daughters converges with similar findings from other mammals, where philopatry facilitates further cooperative or tolerative interactions [11,53]. This includes a growing literature documenting female-biased kin benefits in less social mammalian species. For example, Eurasian lynx (*Lynx lynx*) mothers and daughters appear to share space when resources are patchy [28]. It is worth noting the magnitude of the effect of proximity to daughters in red squirrels. For example, among females whose nearest primary kin was a daughter, having that daughter 30 m away rather than 130 m away corresponded to a change in the mother’s predicted probability of overwinter survival from approximately 37% to 82% (calculated for a 2-year old female in a non-mast year with average dyadic familiarity; table S2). Interestingly, daughters did not appear to receive similar benefits from access to their mother. We also note the lack of any influences of nearby kin on annual reproductive success in females. Social thermoregulation is a possible mechanism driving these kin-based increases in survival, which is known to occur among females whose territories are close together (mean of 43.4±18.0 m SD apart) [24]. However, we would generally expect the benefits of social thermoregulation to be reciprocal, i.e., also increase survival for daughters living near mothers [54]. Previous work has shown that territorial intrusions (possibly to pilfer cones) sometimes involve mothers invading offspring territories, and again is more likely to occur when distances between territories are small [29]. Thus, a seemingly more likely explanation for these unidirectional benefits is that mothers increase their survival by maintaining access to the resources of nearby daughters. It is important to note though that generally, pilfered cones appear to make up a very small proportion of an individual’s food stores in this population [55]. An additional explanation for these patterns is that generally successful mothers are both more likely to recruit daughters nearby and to survive. However, the benefits of access to daughters were detected within the sample of mothers who had a daughter within 130 m – that daughters benefited mothers only when living very close makes it less likely that correlations between reproductive output and survival are the major driver of these patterns.

For male red squirrels, annual survival was not associated with access to kin, but was instead linked to environmental variation (i.e., masting events). This converges with evidence from a separate study which identified a similar, though non-significant, trend of reduced survival during mast years in this population [56]. We hypothesize that this reduction in overwinter survival for males may be a cost of intense investment in reproduction: males expand their territory sizes nearly three-fold during mast years [33], possibly a result of increased efforts to find and mate with receptive females.

Much more surprising was the finding that male red squirrels had increased reproductive success when living closer to their fathers. A typical male living 50 m from its father was predicted to sire 2 pups on average, but no pups if that father lived 130 m away (calculated for 2-year old during non-mast year with average dyadic familiarity). This was unexpected for two reasons: first, while red squirrels appear to discriminate kin and non-kin based on rattle vocalizations [57], it remains unclear whether this recognition stems from shared early life (natal) experience [58], which would limit recognition to maternal kin. This points to the possibility of another mechanism by which fathers and offspring can recognize one another, such as phenotypic matching [58,59], thought to facilitate kin recognition in species of many taxa (e.g., cichlids (e.g., *Pelvicachromis taeniatus*) [60]; Siberian jays (*Perisoreus infaustus*) [61]. Second, male red squirrels have been found to have enhanced lifetime reproductive success as immigrants to a new population [62], congruent with the expectation that they are avoiding competition with related males. In a similar vein, evidence in this study of reduced male reproductive success when closer to female kin (figure 3), could reflect the costs of reduced access to unrelated mates. However, there appears to be little inbreeding avoidance in this population: offspring from related parents are common (*r* > 0.125 for 19% of mates) and do not face reductions in birth mass, growth, or survival [63], suggesting that other factors may explain this pattern.

In some species, males form reproductive alliances to cooperate in securing and defending mating opportunities with females, leading to improved reproductive success [64–67]. Alliance members are often related, leading to indirect benefits for individuals that do not enjoy increased mating opportunities themselves. Much rarer associations between fathers and sons have been linked to reproductive benefits in red howler monkeys (*Alouatta seniculus*) [68] and yellow baboons (*Papio cynocephalus*) [69], for example. And while male-male alliances are typically found in group-living species, it has been suggested that rudimentary aspects of coalitionary behaviour may be exhibited in otherwise solitary males [70]. So, could the observed ARS benefits observed among father-son dyads in red squirrels be evidence of cryptic coalitions? Theory suggests that male reproductive coalitions are more likely to form when 1) females are receptive for short periods of time and 2) cannot be monopolized by a single mate [70,71]. Both are true of the “scramble competition” mating system of red squirrels, where estrus lasts just a single day, during which an estrous female mates with an average of 7 males [36,63]. However, male red squirrels are not known to engage in any observable affiliative interactions, making even basic aspects of coalitionary behaviour unlikely. Perhaps more likely is that fathers and sons are simply less likely to disrupt one another’s copulations during mating chases. Hypothetically, related males living together could also benefit from socially transmitted information about the locations of receptive females. One avenue for future work would be to investigate where proximity between fathers and sons predicts co-parentage of offspring in the same litter (above and beyond any influences of distance to the mother), expected if male kin are somehow facilitating mating opportunities. Alternatively, increased tolerance among fathers and sons could reduce the need for defense and thereby allow males to exert more energy searching for mates.

More generally, these findings place an even greater emphasis on territory acquisition as a determinant of fitness for red squirrels [72], by increasing the stakes of co-locating with certain types of kin. Red squirrels are one of very few species known to exhibit bequeathal, where an adult female undergoes breeding dispersal thus allowing an offspring (with a slight bias for daughters over sons) to inherit her territory and resource cache [21,73]. Such an extreme display of nepotism is striking, given the critical importance of territory ownership for survival. Interestingly then, previous work in this population has not identified a survival cost for bequeathing mothers [74]. Theory suggests that if a mother can secure a new territory near their previous site, increased access to offspring might lead to greater downstream fitness benefits via kin cooperation [75]. Consistent with this hypothesis, our findings suggest that bequeathal may provide additional, direct benefits to mothers in addition to enhancements in offspring survival [74], at least when the inheritor is a daughter. Compared to females, male red squirrels tend to live farther from primary kin, with no obvious pattern of father-son dyads living closer together than father-daughter dyads, for example (see electronic supplement, figure S1). Thus we find no obvious indication of kin-biased dispersal, whereby males might be expected to preferentially acquire territories near male kin [67,76], though a thorough test of this possibility may be a worthwhile direction for future work.

Leveraging geographic distance among kin as a proxy for their ability to interact allowed us to directly measure fitness consequences without a clear sense of the specific behaviours that are important [31]. Beyond age and resource pulses (which were controlled for in our models), we are unaware of any likely confounds of proximity among kin and individual fitness in the population. Moreover, the importance of small differences in distance to kin (i.e., within the 130 m social neighbourhood) and the variation in effects across genealogical relationship are difficult to explain without distance-constrained social behaviours. However, future work is necessary to understand the contribution of behavioural interactions or other processes in causing these patterns.

Consistent with previous findings [23], male squirrels also had enhanced reproductive success when living near familiar individuals. Unlike previous work however, we did not find significant effects of familiarity on survival for either males or females. We expect that this difference is a result of the dyadic approach in the present study – in other words, the benefits of familiarity may be small when considering a single target individual. This distinction highlights the separate spatial and social scales at which familiarity and kinship operate to influence red squirrel fitness. Whereas the benefits of familiarity emerge primarily as a property of a wider socio-spatial environment (the red squirrel neighbourhood) [23], the benefits of kinship occur via close proximity to a small set of specific individuals. Crucially, access to primary kin may not necessarily translate into variation in local genetic relatedness, raising questions about the use of local relatedness as a measure of kinship dynamics (reviewed in [9]).

The idiosyncratic nature of our results suggests that consequences of access to kin may be overlooked in systems where associations are not parsed into specific genealogical relationships. It also suggests that squirrels require kin recognition to maintain these relationships, contrary to many species where spatial heuristics (e.g., “cooperate with those around you”) are sufficient for cooperation [77,78]. In tandem with previous studies, our work contributes to an increasingly complex depiction of sociality in a mostly solitary species. Though further work is needed to understand the specific mechanisms causing these benefits, it appears that kinship can have multifaceted and impactful contributions to fitness in solitary mammals

## Supporting information

Electronic supplement

## Acknowledgements

We thank Agnes MacDonald and her family for long-term access to her trapline, and the Champagne and Aishihik First Nations for allowing us to conduct our work within their traditional territory. We are grateful to each of the field technicians who have contributed to the long-term Kluane Red Squirrel Project database over the years. We thank Erin Siracusa and April Martinig for providing helpful advice regarding analysis and interpretation. We also thank members of the McAdam lab including Shelby Bohn, David Delaney, Simon Denomme-Brown, Alex Hare, Katie Kariatsumari, and Quinn Webber for valuable and supportive commentary on an earlier version of this work. Lastly, we thank two anonymous reviewers for helpful comments that improved this work.

