## Supplementary material for "Benefits of living closer to kin vary by genealogical relationship in a territorial mammal": Electronic supplement

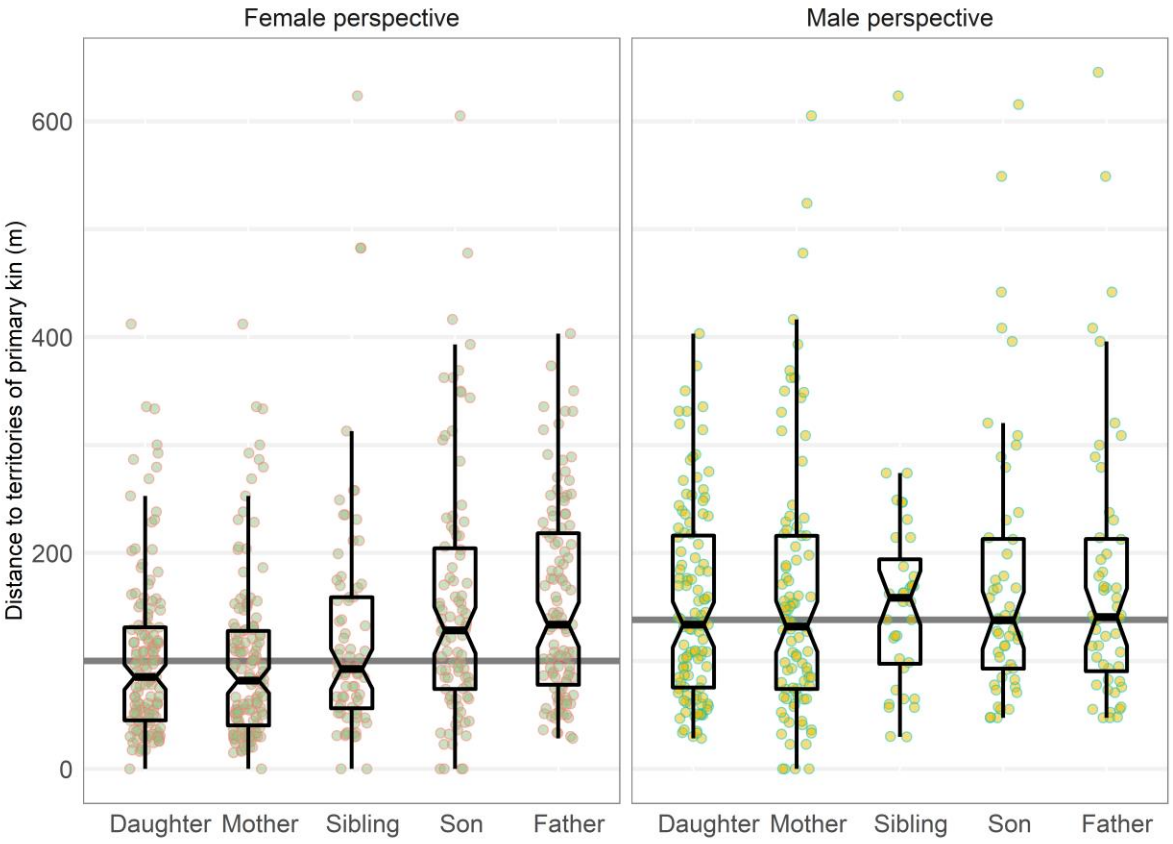


**Figure S1. Distance to primary kin varies by sex and genealogical relationship.** Boxplots show distributions of linear distances between territories of red squirrels included in our study, shown separately for the sex of the focal individual and different types of primary kin. For this figure, values were subset to include just one measure for each directional, focal-target dyad. The grey horizontal line represents the median distance to primary kin, by sex, and is higher for males than females. Note also that some boxes are expected to contain redundant information (e.g., ‘Daughter’ and ‘Mother’ in the left panel).

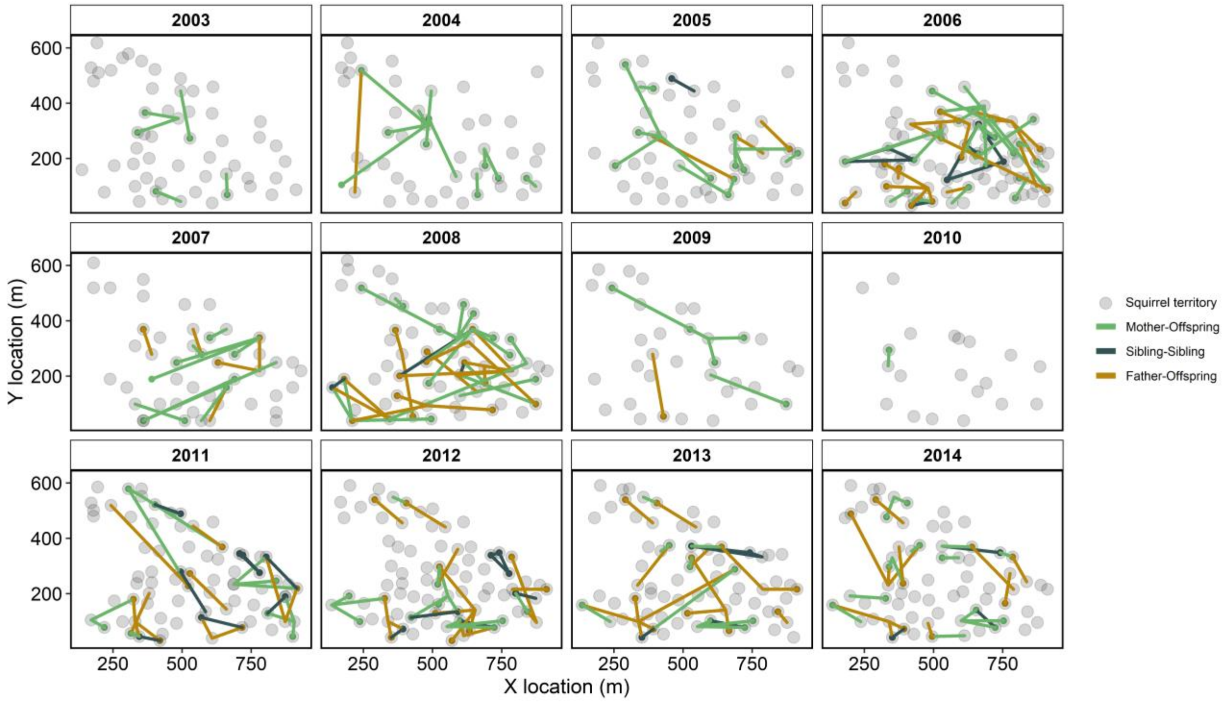


**Figure S2. Spatial overview of kin relationships on "Sulphur" study area.** Lines linking certain individuals represent relationships among primary kin, with colors distinguishing dyads as a sibling pair or by the sex of the parent. Grey circles show the locations of middens or territory centres for all territories occupied each year. Mast years (resource pulses) occurred in 2005, 2010, and 2014.

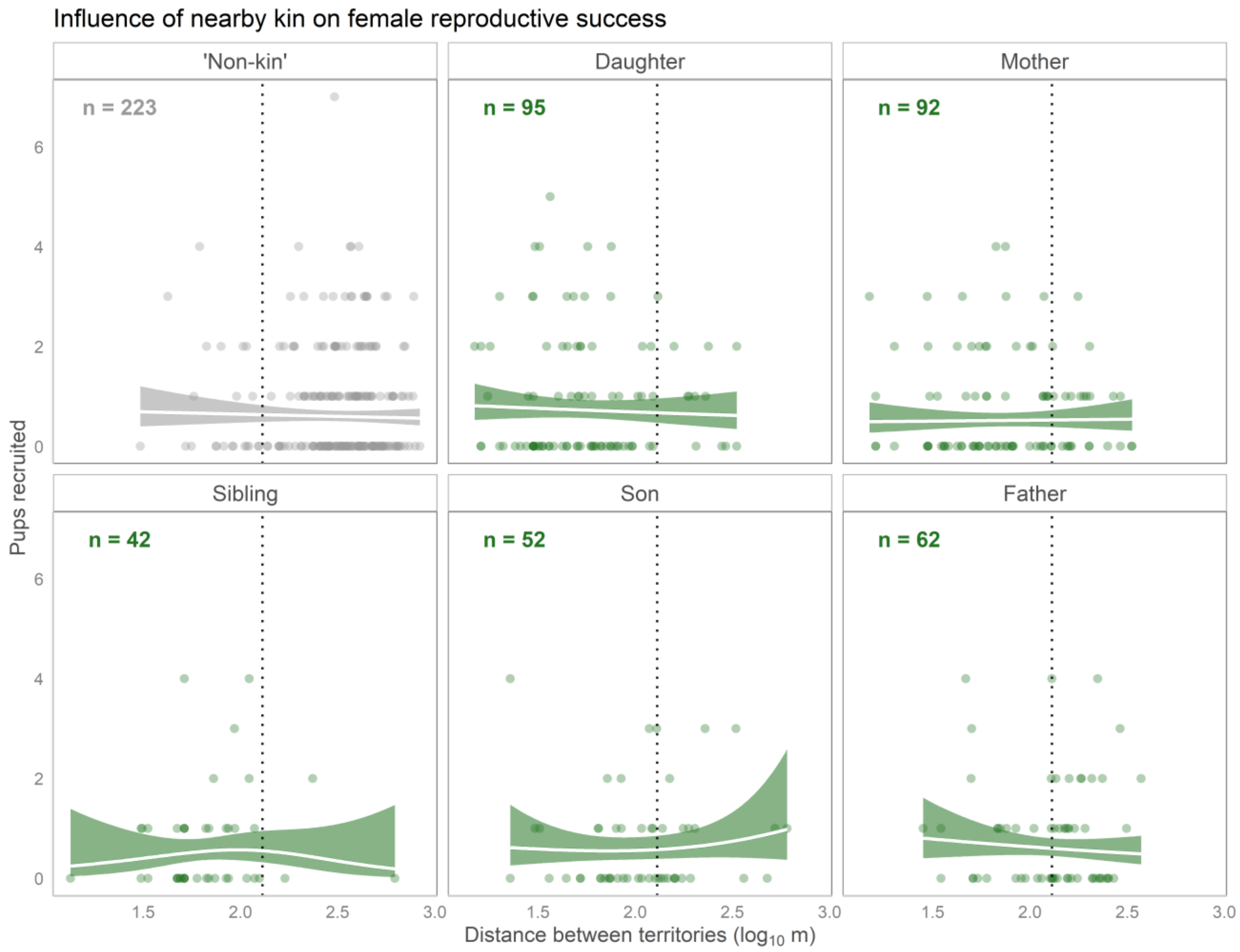


**Figure S3. GAM-generated relationships between female ARS and distance to kin.** Lines represent predicted probabilities of the number of pups a breeding female successfully recruits, estimated with GAMs, controlling for the influences of age and resource pulses, with shaded bands representing 95% confidence intervals. Here, probabilities are shown for a focal individual of age 2 in a non-mast year. Dots represent raw data, so are not an exact reflection of the partial smooth. The dotted vertical line corresponds to 130m, a commonly used radius of red squirrels' social neighbourhoods. Non-kin records are made up of individuals with no primary kin in the study area, and distances were calculated relative to a randomly chosen target individual.

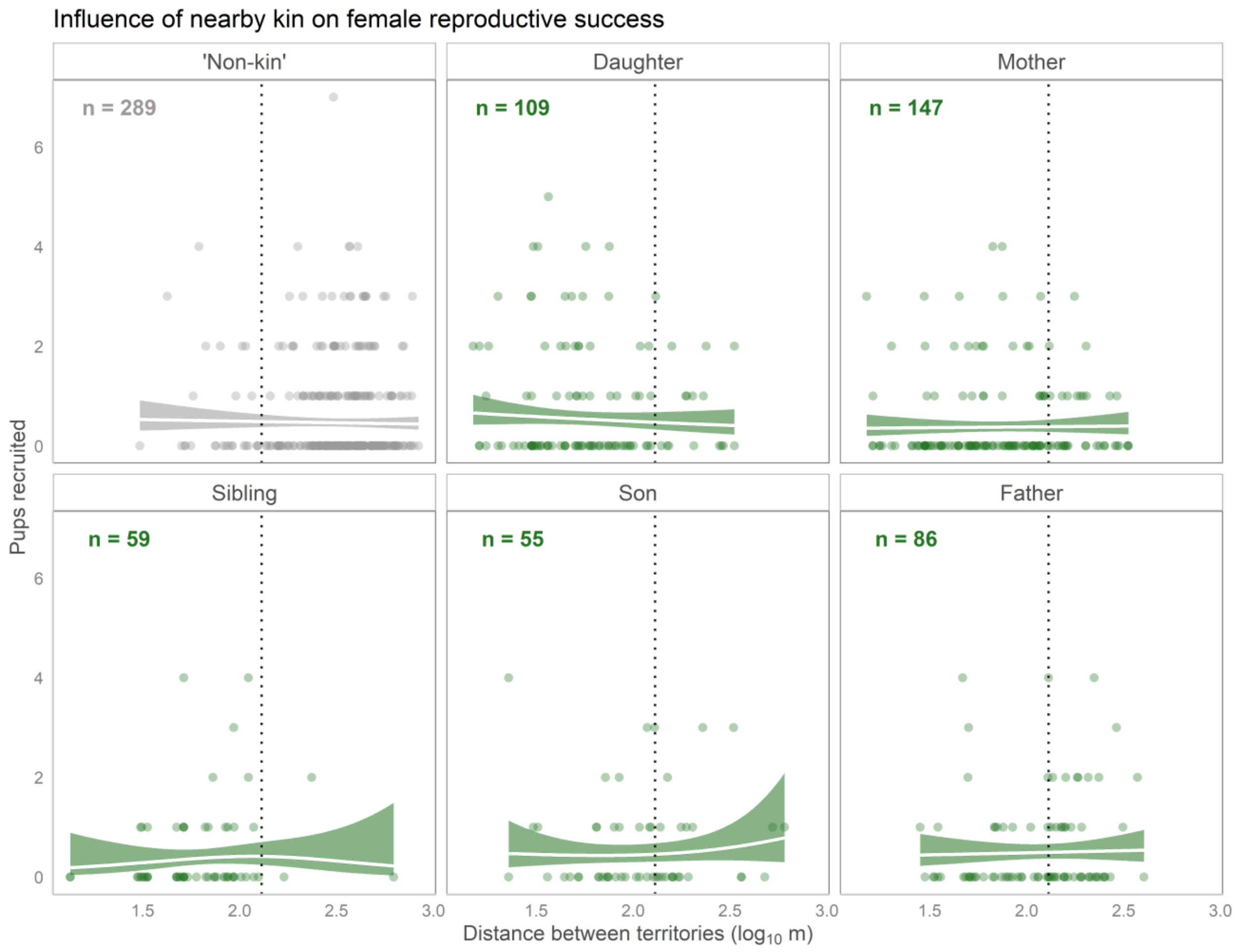


**Figure S4. Inclusion of non-breeders did not alter results for female ARS.** Lines represent predicted probabilities of the number of pups a breeding female successfully recruits, estimated with GAMs, controlling for the influences of age and resource pulses, with shaded bands representing 95% confidence intervals. Dots represent raw data, so are not an exact reflection of the partial smooth. Here, probabilities are shown for a focal individual of age 2 in a non-mast year. The dotted vertical line corresponds to 130m, a commonly used radius of red squirrels' social neighbourhoods. Non-kin records are made up of individuals with no primary kin in the study area, and distances were calculated relative to a randomly chosen target individual.

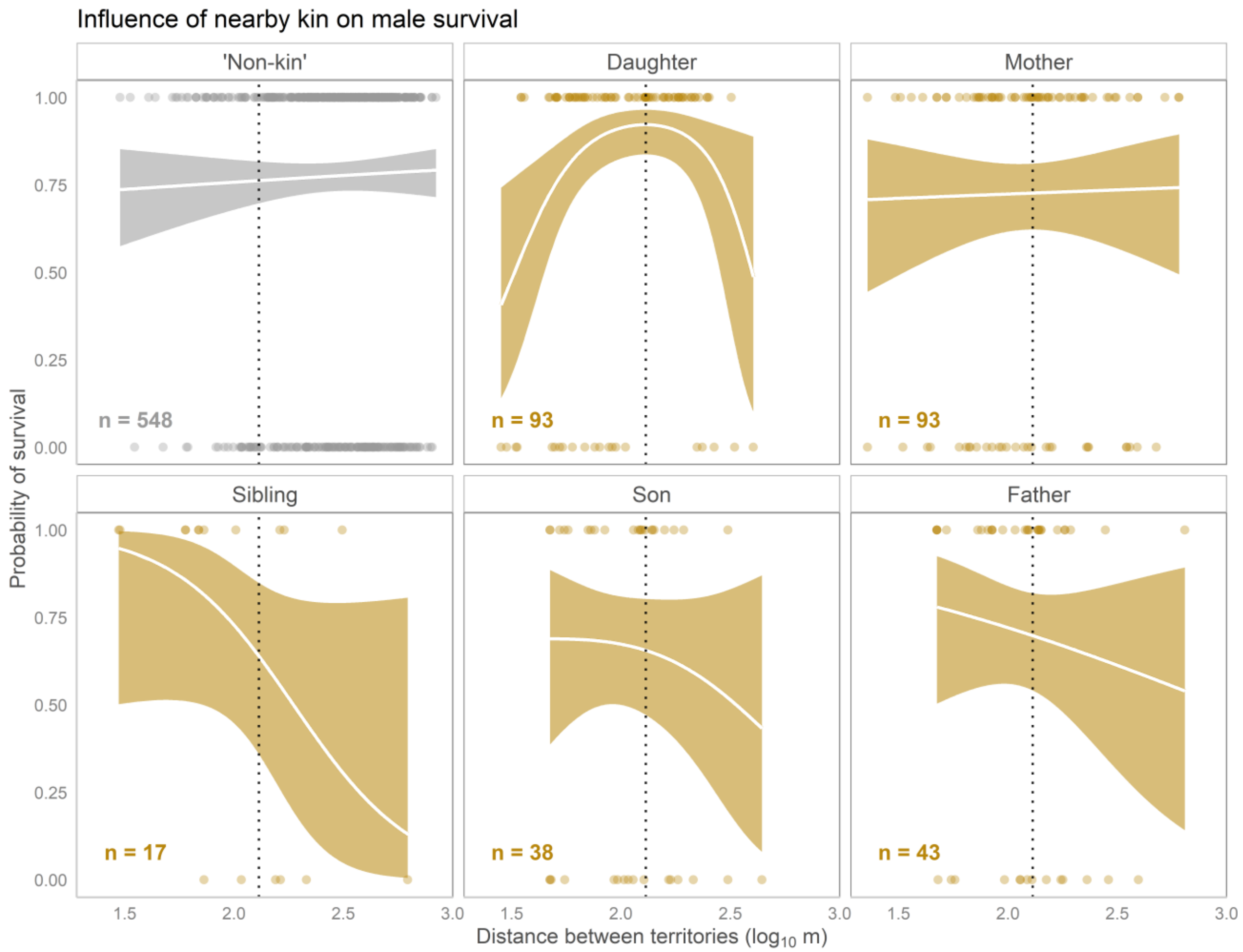


**Figure S5. GAM-generated relationships between male survival and distance to kin.** Lines represent predicted probabilities of male survival, estimated with GAMs, controlling for the influences of age and resource pulses, with shaded bands representing 95% confidence intervals. Dots represent raw data, so are not an exact reflection of the partial smooth. Here, probabilities are shown for a focal individual of age 2 in a non-mast year. The dotted vertical line corresponds to 130m, a commonly used radius of red squirrels' social neighbourhoods. Non-kin records are made up of individuals with no primary kin in the study area, and distances were calculated relative to a randomly chosen target individual.

Secondary models based on the entire study area with quadratic estimates for influences of kin:

We fit additional parametric models including measures from individuals with primary kin at greater distances than the 130m social neighborhood. Because initial GAM visualizations suggested that fitness-distance relationships sometimes shifted when considering the full range of distances in each study area, these were fit with quadratic effects of distance, using the *poly* function in R [1]. Otherwise, GLMM predictors and response variables, structures, transformation decisions, statistical packages used, and diagnostic procedures were identical to those described in the main text for the social neighbourhood models.

Outputs from these models are presented on the following pages (tables S3, S5, S7, S9). As noted in the main text, results from these “full study area” models produced generally similar results to the models based on the social neighbourhood. Effects that differed at the larger spatial scale are highlighted in the main text and again as follows:

- For female annual survival across the entire study area, the effect of daughters was non-significant, suggesting that influences of daughters are only important at the scale of the social environment (130 m neighbourhood).
- For male annual survival across the study area, a significant, negative effect of the quadratic distance-genealogical relationship interaction term for daughters was observed (table S5). This negative parabola was consistent with the shape of the relationship indicated by the positive, non-significant ($\log_{10}$Distance × Daughter) term in the social neighbourhood model (table S4), suggesting that across a large spatial scale, males have the greatest annual reproductive success when daughters live at intermediate distances.
- For female annual reproductive success across the study area (table S7), results were nearly identical to those from the social neighbourhood model (Table S6), suggesting no influence of access to kin on female reproductive success at either spatial scale.
- For male annual reproductive success across the study area (table S9), the distance term for the reference category (non-kin) was negative and significant, suggesting that at a large spatial scale, males sired more pups when living more closely to other squirrels (excluding primary kin). The influence of daughters emerged as a significant influence on the number of pups sired. One point of potential confusion is that the linear terms for daughters and mothers ($\log_{10}$Distance × Daughter) and ($\log_{10}$Distance × Daughter) have

opposite signs, despite having a similarly negative effect on male annual reproductive success in the social neighbourhood model. It is important to simultaneously consider the quadratic terms however, ($\log_{10}\text{Distance}^2$ × Daughter) and ($\log_{10}\text{Distance}^2$ × Mother), which indicate a negative parabolic effect for the influence of daughters across the study grid, and a positive linear effect for the influence of mothers across the study grid, both of which result in a positive link between proximity to daughters and male reproductive success *at close distances* (as reflected in the social neighbourhood model). This pattern is also apparent in figure 2 in the main text.

**Table S1**. Results from generalized linear mixed effects model estimating the influence of sex on distance to nearest primary kin (parent, offspring, full sibling) in North American red squirrels.

| **Sex differences in distance to kin** | | | | | | |
|---|---|---|---|---|---|---|
| | | Variable | Estimate | 95% CI | *z* | *p* |
| $n_{Obs}$ | 346 | Intercept | 4.60 | (4.49, 4.71) | 102.44 | < 0.001 |
| $n_{Focal}$ | 298 | $Sex_{Focal}$: Male | 0.35 | (0.33, 0.56) | 4.26 | < 0.001 |
| | | Focal ID (random effect variance) | 0.18 | | | |

**Table S2**. Results from generalized linear mixed effects model estimating the influences of distance to primary kin on annual survival in female North American red squirrels.

| **Female annual survival** | | | | | | |
|---|---|---|---|---|---|---|
| | | Variable | Estimate | 95% CI | *z* | *p* |
| **Within social neighbourhood** | | | | | | |
| $n_{Obs}$ | 619 | Intercept | -0.37 | (-3.6, 2.85) | -0.23 | 0.821 |
| $n_{Focal}$ | 317 | $\log_{10}$Distance | 0.55 | (-1.09, 2.18) | 0.65 | 0.513 |
| $n_{Dyad}$ | 423 | Daughter | 6.19 | (1.05, 11.33) | 2.36 | 0.018 |
| $R^2_{Marg.}$ | 0.06 | Mother | 0.71 | (-3.75, 5.16) | 0.31 | 0.755 |
| $R^2_{Cond.}$ | 0.14 | Sibling | 1.67 | (-4.95, 8.3) | 0.49 | 0.621 |
| | | Son | 8.37 | (-2.28, 19.02) | 1.54 | 0.124 |
| | | Father | -1.14 | (-8.99, 6.7) | -0.29 | 0.775 |
| | | Familiarity | 0.04 | (-0.19, 0.26) | 0.32 | 0.751 |
| | | Age | 0.44 | (-0.12, 1) | 1.55 | 0.122 |
| | | $Age^2$ | -0.10 | (-0.18, -0.02) | -2.44 | 0.015 |
| | | Mast year (Y) | -0.07 | (-0.91, 0.76) | -0.17 | 0.862 |
| | | $\log_{10}$Distance × Daughter | -3.78 | (-6.66, -0.9) | -2.57 | 0.010 |
| | | $\log_{10}$Distance × Mother | -0.33 | (-2.76, 2.11) | -0.26 | 0.793 |
| | | $\log_{10}$Distance × Sibling | -0.73 | (-4.4, 2.94) | -0.39 | 0.697 |
| | | $\log_{10}$Distance × Son | -4.31 | (-9.88, 1.27) | -1.51 | 0.130 |
| | | $\log_{10}$Distance × Father | 0.76 | (-3.54, 5.07) | 0.35 | 0.728 |
| | | Focal ID (random effect variance) | $2.13 \times 10^{-8}$ | | | |
| | | Dyad ID (random effect variance) | 0.188 | | | |
| | | Year (random effect variance) | 0.264 | | | |

**Table S3**. Results from generalized linear mixed effects model estimating the influences of distance to primary kin on annual survival in female North American red squirrels.

| **Female annual survival: full study area model** | | | | | | |
|---|---|---|---|---|---|---|
| | | Variable | Estimate | 95% CI | *z* | *p* |
| **Across entire study area** | | | | | | |
| $n_{Obs}$ | 745 | Intercept | 0.58 | (-0.28, 1.44) | 1.32 | 0.185 |
| $n_{Focal}$ | 349 | $\log_{10}$Distance | 6.75 | (-9.94, 23.45) | 0.79 | 0.428 |
| $n_{Dyad}$ | 533 | $\log_{10}$Distance$^2$ | 3.19 | (-8.7, 15.09) | 0.53 | 0.599 |
| $R^2_{Marg.}$ | 0.07 | Daughter | -0.16 | (-1.12, 0.81) | -0.32 | 0.748 |
| $R^2_{Cond.}$ | 0.12 | Mother | 0.32 | (-0.48, 1.13) | 0.79 | 0.431 |
| | | Sibling | 1.69 | (-1.41, 4.78) | 1.07 | 0.286 |
| | | Son | 0.28 | (-0.64, 1.2) | 0.59 | 0.552 |
| | | Father | 0.42 | (-0.6, 1.45) | 0.81 | 0.420 |
| | | Familiarity | 0.05 | (-0.15, 0.26) | 0.48 | 0.626 |
| | | Age | 0.36 | (-0.13, 0.84) | 1.44 | 0.150 |
| | | Age$^2$ | -0.09 | (-0.16, -0.02) | -2.58 | 0.010 |
| | | Mast year (Y) | -0.17 | (-0.83, 0.5) | 0.49 | 0.628 |
| | | $\log_{10}$Distance × Daughter | -9.51 | (-38.52, 19.49) | 1.44 | 0.520 |
| | | $\log_{10}$Distance × Mother | -2.62 | (-28.76, 23.52) | -2.58 | 0.844 |
| | | $\log_{10}$Distance × Sibling | 27.40 | (-62.75, 117.55) | -0.48 | 0.551 |
| | | $\log_{10}$Distance × Son | -24.11 | (-53.35, 5.12) | -0.64 | 0.106 |
| | | $\log_{10}$Distance × Father | -0.26 | (-30.35, 29.84) | -0.20 | 0.987 |
| | | $\log_{10}$Distance$^2$ × Daughter | 12.14 | (-10.75, 35.03) | 0.60 | 0.298 |
| | | $\log_{10}$Distance$^2$ × Mother | 0.32 | (-19.1, 19.74) | -1.62 | 0.974 |
| | | $\log_{10}$Distance$^2$ × Sibling | 15.17 | (-38.27, 68.62) | -0.02 | 0.578 |
| | | $\log_{10}$Distance$^2$ × Son | -1.82 | (-28.11, 24.47) | 1.04 | 0.892 |
| | | $\log_{10}$Distance$^2$ × Father | -3.30 | (-37.4, 30.79) | 0.03 | 0.849 |
| | | Focal ID (random effect variance) | $9.98 \times 10^{-5}$ | | | |
| | | Dyad ID (random effect variance) | 0.101 | | | |
| | | Year (random effect variance) | 0.150 | | | |

**Table S4**. Results from generalized linear mixed effects model estimating the influences of distance to primary kin on annual survival in male North American red squirrels.

| **Male annual survival** | | | | | | |
|---|---|---|---|---|---|---|
| | | Variable | Estimate | 95% CI | $z$ | $p$ |
| **Within social neighbourhood** | | | | | | |
| $n_{Obs}$ | 710 | Intercept | 0.74 | (-1.61, 3.1) | 0.62 | 0.536 |
| $n_{Focal}$ | 354 | $log_{10}$Distance | 0.30 | (-0.86, 1.46) | 0.51 | 0.607 |
| $n_{Dyad}$ | 570 | Daughter | -5.27 | (-12.3, 1.75) | -1.47 | 0.141 |
| $R^2_{Marg.}$ | 0.07 | Mother | 0.38 | (-5.66, 6.43) | 0.12 | 0.901 |
| $R^2_{Cond.}$ | 0.10 | Sibling | 20.13 | (-15.36, 55.61) | 1.11 | 0.266 |
| | | Son | -1.04 | (-10.93, 8.84) | -0.21 | 0.836 |
| | | Father | 3.34 | (-7.49, 14.16) | 0.60 | 0.546 |
| | | Familiarity | 0.11 | (-0.09, 0.3) | 1.06 | 0.288 |
| | | Age | 0.12 | (-0.38, 0.62) | 0.45 | 0.650 |
| | | $Age^2$ | -0.06 | (-0.14, 0.01) | -1.65 | 0.098 |
| | | Mast year (Y) | -0.69 | (-1.33, -0.05) | -2.10 | 0.036 |
| | | $log_{10}$Distance × Daughter | 3.00 | (-0.88, 6.88) | 1.52 | 0.129 |
| | | $log_{10}$Distance × Mother | -0.51 | (-3.74, 2.71) | -0.31 | 0.755 |
| | | $log_{10}$Distance × Sibling | -10.68 | (-29.21, 7.86) | -1.13 | 0.259 |
| | | $log_{10}$Distance × Son | 0.25 | (-4.93, 5.43) | 0.09 | 0.925 |
| | | $log_{10}$Distance × Father | -2.05 | (-7.66, 3.56) | -0.71 | 0.475 |
| | | Focal ID (random effect variance) | $3.81 \times 10^{-5}$ | | | |
| | | Dyad ID (random effect variance) | $2.38 \times 10^{-5}$ | | | |
| | | Year (random effect variance) | 0.133 | | | |

**Table S5**. Results from generalized linear mixed effects model estimating the influences of distance to primary kin on annual survival in male North American red squirrels.

| **Male annual survival: full study area model** | | | | | | |
|---|---|---|---|---|---|---|
| | | Variable | Estimate | 95% CI | *z* | *p* |
| **Across entire study area** | | | | | | |
| $n_{Obs}$ | 832 | Intercept | 0.88 | (0.21, 1.55) | 2.57 | 0.010 |
| $n_{Focal}$ | 387 | $log_{10}$Distance | -1.63 | (-10.3, 7.05) | -0.37 | 0.713 |
| $n_{Dyad}$ | 717 | $log_{10}$Distance$^2$ | -0.01 | (-7.71, 7.69) | 0.00 | 0.998 |
| $R^2_{Marg.}$ | 0.08 | Daughter | -0.35 | (-1.46, 0.75) | -0.63 | 0.531 |
| $R^2_{Cond.}$ | 0.10 | Mother | -0.29 | (-0.94, 0.35) | -0.88 | 0.377 |
| | | Sibling | -1.19 | (-2.68, 0.3) | -1.56 | 0.118 |
| | | Son | -1.57 | (-3.19, 0.05) | -1.89 | 0.058 |
| | | Father | -0.57 | (-1.61, 0.47) | -1.08 | 0.281 |
| | | Familiarity | -0.03 | (-0.21, 0.16) | -0.28 | 0.776 |
| | | Age | 0.41 | (-0.05, 0.87) | 1.76 | 0.078 |
| | | Age$^2$ | -0.1 | (-0.17, -0.03) | -2.76 | 0.006 |
| | | Mast year (Y) | -0.62 | (-1.19, -0.05) | -2.14 | 0.032 |
| | | $log_{10}$Distance × Daughter | -26.55 | (-60.35, 7.24) | -1.54 | 0.124 |
| | | $log_{10}$Distance × Mother | 1.62 | (-17.18, 20.42) | 0.17 | 0.866 |
| | | $log_{10}$Distance × Sibling | -28.60 | (-71.35, 14.14) | -1.31 | 0.190 |
| | | $log_{10}$Distance × Son | -25.06 | (-70.55, 20.43) | -1.08 | 0.280 |
| | | $log_{10}$Distance × Father | -5.98 | (-35.08, 23.13) | -0.40 | 0.687 |
| | | $log_{10}$Distance$^2$ × Daughter | -30.31 | (-53.52, -7.1) | -2.56 | 0.010 |
| | | $log_{10}$Distance$^2$ × Mother | -1.36 | (-15.37, 12.64) | -0.19 | 0.849 |
| | | $log_{10}$Distance$^2$ × Sibling | 13.16 | (-26.79, 53.11) | 0.65 | 0.519 |
| | | $log_{10}$Distance$^2$ × Son | -20.07 | (-54.61, 14.46) | -1.14 | 0.255 |
| | | $log_{10}$Distance$^2$ × Father | 2.35 | (-22.27, 26.98) | 0.19 | 0.851 |
| | | Focal ID (random effect variance) | $3.03 \times 10^{-5}$ | | | |
| | | Dyad ID (random effect variance) | $6.14 \times 10^{-5}$ | | | |
| | | Year (random effect variance) | 0.099 | | | |

**Table S6**. Results from generalized linear mixed effects model estimating the influences of distance to primary kin on annual reproductive success (number of pups successfully recruited) in female North American red squirrels.

| **Female annual reproductive success** | | | | | | |
|---|---|---|---|---|---|---|
| | | Variable | Estimate | 95% CI | *z* | *p* |
| **Within social neighbourhood** | | | | | | |
| $n_{Obs}$ | 465 | Intercept | -2.38 | (-4.39, -0.38) | -2.33 | 0.020 |
| $n_{Focal}$ | 259 | $log_{10}$Distance | 0.86 | (-0.14, 1.85) | 1.68 | 0.092 |
| $n_{Dyad}$ | 342 | Daughter | 2.06 | (-0.53, 4.64) | 1.56 | 0.118 |
| $R^2_{Marg.}$ | 0.17 | Mother | 1.01 | (-1.79, 3.82) | 0.71 | 0.480 |
| $R^2_{Cond.}$ | 0.27 | Sibling | 0.40 | (-4.26, 5.05) | 0.17 | 0.867 |
| | | Son | 1.84 | (-2.06, 5.75) | 0.92 | 0.355 |
| | | Father | 3.2 | (-1.42, 7.82) | 1.36 | 0.175 |
| | | Familiarity | 0.04 | (-0.07, 0.15) | 0.72 | 0.471 |
| | | Age | 0.11 | (-0.21, 0.43) | 0.66 | 0.507 |
| | | $Age^2$ | -0.02 | (-0.07, 0.02) | -0.95 | 0.341 |
| | | Mast year (Y) | 1.16 | (0.81, 1.52) | 6.40 | < 0.001 |
| | | $log_{10}$Distance × Daughter | -1.03 | (-2.46, 0.41) | -1.40 | 0.160 |
| | | $log_{10}$Distance × Mother | -0.58 | (-2.1, 0.94) | -0.75 | 0.456 |
| | | $log_{10}$Distance × Sibling | -0.26 | (-2.76, 2.23) | -0.21 | 0.836 |
| | | $log_{10}$Distance × Son | -1.03 | (-3.1, 1.04) | -0.97 | 0.330 |
| | | $log_{10}$Distance × Father | -1.67 | (-4.14, 0.8) | -1.32 | 0.186 |
| | | Focal ID (random effect variance) | 0.062 | | | |
| | | Dyad ID (random effect variance) | 0.043 | | | |
| | | Year (random effect variance) | 0.046 | | | |

**Table S7**. Results from generalized linear mixed effects model estimating the influences of distance to primary kin on annual reproductive success (number of pups successfully recruited) in female North American red squirrels.

| **Female annual reproductive success: full study area model** | | | | | | |
|---|---|---|---|---|---|---|
| | | Variable | Estimate | 95% CI | *z* | *p* |
| **Across entire study area** | | | | | | |
| $n_{Obs}$ | 565 | Intercept | -1.45 | (-3, 0.09) | -1.85 | 0.064 |
| $n_{Focal}$ | 293 | $log_{10}$Distance | 0.35 | (-0.26, 0.95) | 1.13 | 0.260 |
| $n_{Dyad}$ | 427 | Daughter | 1.13 | (-0.82, 3.08) | 1.14 | 0.255 |
| $R^2_{Marg.}$ | 0.16 | Mother | 0.30 | (-1.8, 2.41) | 0.28 | 0.778 |
| $R^2_{Cond.}$ | 0.25 | Sibling | 1.16 | (-1.85, 4.16) | 0.75 | 0.450 |
| | | Son | -0.03 | (-2.82, 2.76) | -0.02 | 0.984 |
| | | Father | 1.17 | (-1.62, 3.96) | 0.82 | 0.411 |
| | | Familiarity | 0.08 | (-0.01, 0.17) | 1.67 | 0.094 |
| | | Age | 0.07 | (-0.21, 0.35) | 0.48 | 0.633 |
| | | $Age^2$ | -0.02 | (-0.06, 0.02) | -1.04 | 0.299 |
| | | Mast year (Y) | 1.12 | (0.73, 1.51) | 5.63 | < 0.001 |
| | | $log_{10}$Distance × Daughter | -0.42 | (-1.33, 0.5) | -0.9 | 0.370 |
| | | $log_{10}$Distance × Mother | -0.15 | (-1.14, 0.83) | -0.31 | 0.759 |
| | | $log_{10}$Distance × Sibling | -0.65 | (-2.14, 0.84) | -0.85 | 0.395 |
| | | $log_{10}$Distance × Son | 0.06 | (-1.2, 1.32) | 0.09 | 0.926 |
| | | $log_{10}$Distance × Father | -0.52 | (-1.78, 0.73) | -0.82 | 0.414 |
| | | Focal ID (random effect variance) | 0.060 | | | |
| | | Dyad ID (random effect variance) | 0.019 | | | |
| | | Year (random effect variance) | 0.065 | | | |

**Table S8**. Results from generalized linear mixed effects model estimating the influences of distance to primary kin on annual reproductive success (number of pups sired) in male North American red squirrels.

| **Male annual reproductive success** | | | | | | |
|---|---|---|---|---|---|---|
| | | Variable | Estimate | 95% CI | $z$ | $p$ |
| **Within social neighbourhood** | | | | | | |
| $n_{Obs}$ | 701 | Intercept | -0.59 | (-1.96, 0.78) | -0.85 | 0.397 |
| $n_{Focal}$ | 346 | $\log_{10}$Distance | -0.1 | (-0.77, 0.56) | -0.31 | 0.757 |
| $n_{Dyad}$ | 562 | Daughter | -1.42 | (-5.12, 2.27) | -0.75 | 0.451 |
| $R^2_{Marg.}$ | 0.16 | Mother | -5.79 | (-11.12, -0.47) | -2.13 | 0.033 |
| $R^2_{Cond.}$ | 0.21 | Sibling | -4.30 | (-17.22, 8.63) | -0.65 | 0.515 |
| | | Son | 3.40 | (-1.85, 8.66) | 1.27 | 0.204 |
| | | Father | 9.09 | (0.93, 17.25) | 2.18 | 0.029 |
| | | Familiarity | 0.12 | (0.02, 0.23) | 2.3 | 0.022 |
| | | Age | 0.74 | (0.42, 1.07) | 4.43 | < 0.001 |
| | | $Age^2$ | -0.11 | (-0.16, -0.06) | -4.37 | < 0.001 |
| | | Mast year (Y) | 0.86 | (0.56, 1.16) | 5.63 | < 0.001 |
| | | $\log_{10}$Distance × Daughter | 1.03 | (-0.95, 3.01) | 1.02 | 0.308 |
| | | $\log_{10}$Distance × Mother | 2.79 | (0.01, 5.58) | 1.97 | 0.049 |
| | | $\log_{10}$Distance × Sibling | 2.20 | (-4.84, 9.24) | 0.61 | 0.541 |
| | | $\log_{10}$Distance × Son | -1.65 | (-4.42, 1.13) | -1.16 | 0.245 |
| | | $\log_{10}$Distance × Father | -5.07 | (-9.5, -0.63) | -2.24 | 0.025 |
| | | Focal ID (random effect variance) | 0.117 | | | |
| | | Dyad ID (random effect variance) | $5.06 \times 10^{-9}$ | | | |
| | | Year (random effect variance) | 0.018 | | | |

**Table S9**. Results from generalized linear mixed effects model estimating the influences of distance to primary kin on annual reproductive success (number of pups sired) in male North American red squirrels.

| **Male annual reproductive success: full study area model** | | | | | | |
|---|---|---|---|---|---|---|
| | | Variable | Estimate | 95% CI | *z* | *p* |
| **Across entire study area** | | | | | | |
| $n_{Obs}$ | 822 | Intercept | -1.44 | (-1.86, -1.01) | -6.67 | < 0.001 |
| $n_{Focal}$ | 378 | $log_{10}$Distance | -4.90 | (-9.5, -0.29) | -2.08 | 0.037 |
| $n_{Dyad}$ | 707 | $log_{10}Distance^2$ | -1.96 | (-6.26, 2.34) | -0.89 | 0.373 |
| $R^2_{Marg.}$ | 0.28 | Daughter | -0.79 | (-1.46, -0.13) | -2.34 | 0.019 |
| $R^2_{Cond.}$ | 0.68 | Mother | -0.10 | (-0.52, 0.32) | -0.48 | 0.630 |
| | | Sibling | 0.06 | (-0.78, 0.91) | 0.15 | 0.885 |
| | | Son | -0.33 | (-1.26, 0.6) | -0.7 | 0.487 |
| | | Father | -0.14 | (-0.75, 0.47) | -0.46 | 0.644 |
| | | Familiarity | 0.08 | (0, 0.15) | 2.06 | 0.039 |
| | | Age | 1.09 | (0.8, 1.38) | 7.47 | < 0.001 |
| | | $Age^2$ | -0.16 | (-0.21, -0.12) | -7.12 | < 0.001 |
| | | Mast year (Y) | 0.89 | (0.62, 1.16) | 6.52 | < 0.001 |
| | | $log_{10}$Distance × Daughter | -21.07 | (-39.79, -2.35) | -2.21 | 0.027 |
| | | $log_{10}$Distance × Mother | 13.44 | (2.03, 24.85) | 2.31 | 0.021 |
| | | $log_{10}$Distance × Sibling | 19.41 | (-0.16, 38.99) | 1.94 | 0.052 |
| | | $log_{10}$Distance × Son | -1.93 | (-27.32, 23.45) | -0.15 | 0.881 |
| | | $log_{10}$Distance × Father | 6.83 | (-8.9, 22.55) | 0.85 | 0.395 |
| | | $log_{10}Distance^2$ × Daughter | -13.69 | (-25.7, -1.68) | -2.23 | 0.025 |
| | | $log_{10}Distance^2$ × Mother | -8.26 | (-19.97, 3.45) | -1.38 | 0.167 |
| | | $log_{10}Distance^2$ × Sibling | 6.67 | (-10.25, 23.6) | 0.77 | 0.440 |
| | | $log_{10}Distance^2$ × Son | 5.6 | (-12.93, 24.12) | 0.59 | 0.554 |
| | | $log_{10}Distance^2$ × Father | 14.85 | (1.14, 28.55) | 2.12 | 0.034 |
| | | Focal ID (random effect variance) | 0.352 | | | |
| | | Dyad ID (random effect variance) | 0.372 | | | |
| | | Year (random effect variance) | 0.020 | | | |

### References for Electronic Supplement

1. R Core Development Team. 2021 *R: A language and environment for statistical computing.* Vienna, Austria: R Foundation for Statistical Computing.
